# Moderate developmental alcohol exposure reduces repetitive alternation in a zebrafish model of fetal alcohol spectrum disorders

**DOI:** 10.1101/370072

**Authors:** Madeleine Cleal, Matthew O. Parker

**Affiliations:** School of Pharmacy and Biomedical Science, University of Portsmouth, UK

**Keywords:** prenatal alcohol exposure, fetal alcohol spectrum disorders, repetitive alternation, learning, zebrafish, y-maze

## Abstract

The damaging effects of alcohol on a developing fetus are well known and cause a range of conditions known as fetal alcohol spectrum disorder (FASD). High levels of alcohol exposure lead to physical deformity and severe cognitive deficits, but more moderate exposure leads to a range of subtle cognitive effects such as reduced social behavior, higher propensity to develop addictions, and reduced spatial working memory. Previous studies have demonstrated that following exposure to relatively low levels of ethanol during early brain development (equivalent in humans to moderate exposure) zebrafish display a range of social and behavioral differences. Here, our aim was to test the hypothesis that moderate developmental ethanol exposure would affect aspects of learning and memory in zebrafish. In order to do this, we exposed zebrafish embryos to 20mM [0.12% v/v] ethanol from 2 to 9 dpf to model the effects of moderate prenatal ethanol (MPE) exposure. At 3 months old, adult fish were tested for appetitive and aversive learning, and for spatial alternation in a novel unconditioned y-maze protocol. We found that MPE did not affect appetitive or aversive learning, but exposed-fish showed a robust reduction in repetitive alternations in the y-maze when compared to age matched controls. This study confirms that moderate levels of ethanol exposure to developing embryos have subtle effects on spatial working memory in adulthood. Our data thus suggest that zebrafish may be a promising model system for studying the effects of alcohol on learning and decision-making, but also for developing treatments and interventions to reduce the negative effects of prenatal alcohol.

## Introduction

Consumption of alcohol by women during pregnancy can result in a range of physical and behavioral abnormalities in the fetus, symptoms which are collectively known as fetal alcohol spectrum disorders (FASDs) (1,2). The most severe and easily diagnosed disorder is fetal alcohol syndrome (FAS), which is characterised by craniofacial malformations, central nervous system dysfunction, growth retardation and reduced intellectual abilities (3,4). Although FAS is an extreme case caused by high levels of chronic alcohol abuse, lower levels of alcohol intake have also been shown to cause a range of milder, less obvious symptoms including deficits in social behaviour (5), decision-making and planning (6,7) and an increased susceptibility to substance abuse in later life, even following adoption (i.e., controlling for environmental effects (8)).

Though heavy chronic abuse of alcohol by a pregnant woman leads to obvious symptoms in the child, behavioural symptoms of milder cases of FASD are rarely accompanied by physical deformities and are thus problematic to diagnose (9). As a result, the number of children affected by milder forms of FASDs is likely to be much higher than those reported (10,11). In the absence of physical symptoms, details of any alcohol consumption must derive from self-report and is open to response biases (12). Thus, to better understand the effects of amount, frequency and timing of exposure of the embryo to alcohol, animal models have been used to bridge the gap (13). Traditionally most animal models of FASDs have been carried out in rodents. However, recently zebrafish have come to light as an alternative model for neurobehavioral research, striking a balance between similarities with human and rodent models, complex behavioural interactions, ease of genetic manipulation, low cost of maintenance and high throughput (14–16).

In rodents, the effects of moderate prenatal ethanol exposure on learning have been mixed and unclear, with some conflicting reports of effects on some aspects of learning (13,17,18). This lack of consistency may be due to the complexities associated with rodent models of prenatal exposure, such as dosing regime (injection vs gavage vs voluntary drinking), maternal effects (ie during gestation) and effects of rearing (e.g., cross-fostering vs maternal rearing (19,20). It is critical, therefore, to get a more developed understanding on the effects of moderate exposure to ethanol during early brain development, and zebrafish may offer a useful complementary model organism in which to achieve this.

Since the pioneering work from the Gerlai (5) and Carvan III (18) groups, zebrafish have been proving to be excellent models for examining the effects of low-to-moderate doses of ethanol exposure on the developing embryo on behavioural endpoints. For example, we have shown that zebrafish exposed to moderate developmental alcohol exposure display alterations in adulthood of social and anxiety behaviour, an increased propensity to develop habits, and this corresponded to changes in mRNA expression of genes typically associated with the reward pathway, including dopamine, serotonin, u-opioid and nicotinic acetylcholine receptors (21,22). Despite some evidence that exposed embryos show reductions in ability to learn a spatial two-choice guessing task (18) no studies have carried out a full assessment of the effects of moderate exposure to ethanol during early brain development on different aspects of learning and memory. The aim of this paper was therefore to characterize aspects of learning and memory in adult zebrafish that have been exposed to moderate levels of ethanol during early brain development. We approached our aim by examining appetitive and aversive learning, and repetitive alternation in a novel unconditioned search protocol using a y-maze. Y-maze tests are widely used to measure exploratory behavior in rodents (23–25) and spatial memory in zebrafish (26).

## Methods

### Subjects and ethanol treatment

Embryos (AB wild-type strain) were collected from multiple individual pairings, sorted and cleaned, and placed in groups of ~40/petri dish in a translucent incubator (28°C) on a 14/10- hour light/dark cycle. Ethanol treatment of embryos was as previously described (27,28). Briefly, at 48 hours post-fertilization, embryos were visually inspected and sorted to ensure all were at the same developmental stage (long-pec phase), then transferred to either 20 mM [0.12 percent (v/v), equating to ~0.04 g/dl BAC] ethanol in aquarium water (ethanol-treatment), or to fresh aquarium water with no alcohol (control). At 5-days-post fertilization, they were transferred, still in their treatment medium, into larger containers (10 × 10 × 20 cm [depth × width × length]) containing 500 ml solution (ethanol or aquarium water), and remained in the incubator. During treatment, water/ethanol media were changed daily. Fish remained in the treatment solution for 7 days, until 9-days post fertilization, after which all fish were transferred into fresh aquarium water and placed on our re-circulating system, initially in groups of 40 in 1.4L tanks (Aquaneering Inc., San Diego, CA, USA). Juvenile zebrafish (at 1-month of age) moved to groups of ~20 in 2.8L tanks on the re-circulating system, on a 14/10-hour light/dark cycle, at ~28.5°C. Fish were tested on behavioral procedures at 3 months of age. Fish were fed a mixture of live brine shrimp and flake food 3 times/day (once/day at weekend). No fish was used for multiple protocols, and following the experiment, all ethanol-exposed fish were euthanized (Aqua-Sed™, Vetark, Winchester, UK). Finally, we used a mixture of male and female fish for all behavioral testing. Previous research with zebrafish has not revealed sex effects for developmental alcohol exposure, and sex was not evaluated as a variable in this study.

### Ethical Statement

All experiments were carried out following scrutiny by the University of Portsmouth Animal Welfare and Ethical Review Board, and under license from the UK Home Office (Animals (Scientific Procedures) Act, 1986) [PPL: P9D87106F].

### Materials

Behavioral testing of adults was carried out using the Zantiks (Zantiks Ltd., Cambridge, UK) AD system (https://www.zantiks.com/products/zantiks-ad), a commercially available, fully integrated behavioral testing environment for adult zebrafish ((29), and see Figure 1). All tank inserts were acrylic, with opaque sides and inserts and a transparent base. The test tanks were placed into the Zantiks AD system (see Figure 1a and b) one tank/time. Each Zantiks AD system was fully controlled via any mobile/web enabled device. Figure 1b and c display the tanks used to measure appetitive learning. This Zantiks AD unit is designed to carry out multiple learning protocols in zebrafish but in the present experiment, fish were trained to swim into an initiator zone (Figure 1c, red area) in order to receive a food reinforcer (ZM200 zebrafish food) in the food delivery zone (Figure 1c, orange area). Tanks were filled with 3L water during testing. The total length of the testing environment was 200mm, and 140mm width.

**Figure 1.**
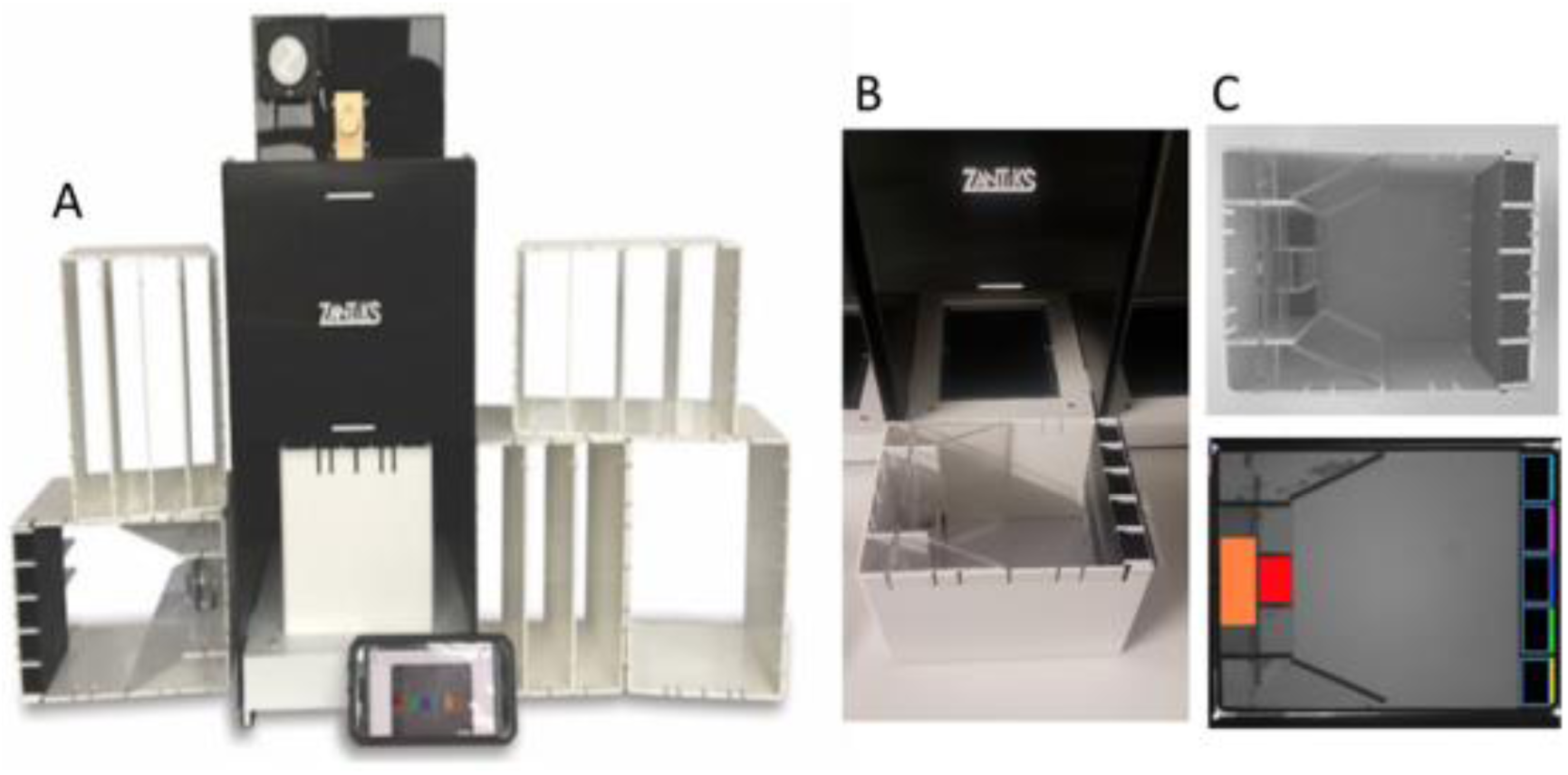
*A)* displays the Zantiks AD testing system used for behavioural testing of adult zebrafish. This is a completely automated system with built in computer for image projection, camera for live imaging and a feeder mechanism. This allows for minimum human disturbance during trials. *1B)* shows the tanks used to measure appetitive learning. *1C)*(top) shows the 5 choice inserts, these can be fitted into each tank and were used in the appetitive learning task. *1C)*(bottom) shows the trigger areas for fish during the 5 choice task. The initiator light (red*) would come on first, fish have to swim into the light to initiate the trial. Following initiation a food reinforce (ZM300 zebrafish food) was delivered into the feeding zone (orange*).When the fish passing through the feeding zone the light is turned off and the initiator light turns back on ready for the next trial. Tanks are filled with 3L water during testing. *In trial lights were all white, colours have only been added for the benefit of the diagram.

Figure 2 displays the tanks equipment used for Pavlovian fear conditioning. The stimuli used were based on previous work (29,30), and comprised either a checker board design (‘check’) (black/white alternating squares) or a dark grey (‘grey’) background. Each tank comprised four lanes (length × width = 160mm × 32mm), separated by opaque acrylic dividers. At each end of the tank was located a steel plate, capable of passing a mild electric current through the tank (9v). Tanks were filled with 1L of water, with ~40mm water at the base. Pilot studies found this amount of water to be optimal for both tracking and conditioning.

**Figure 2.**
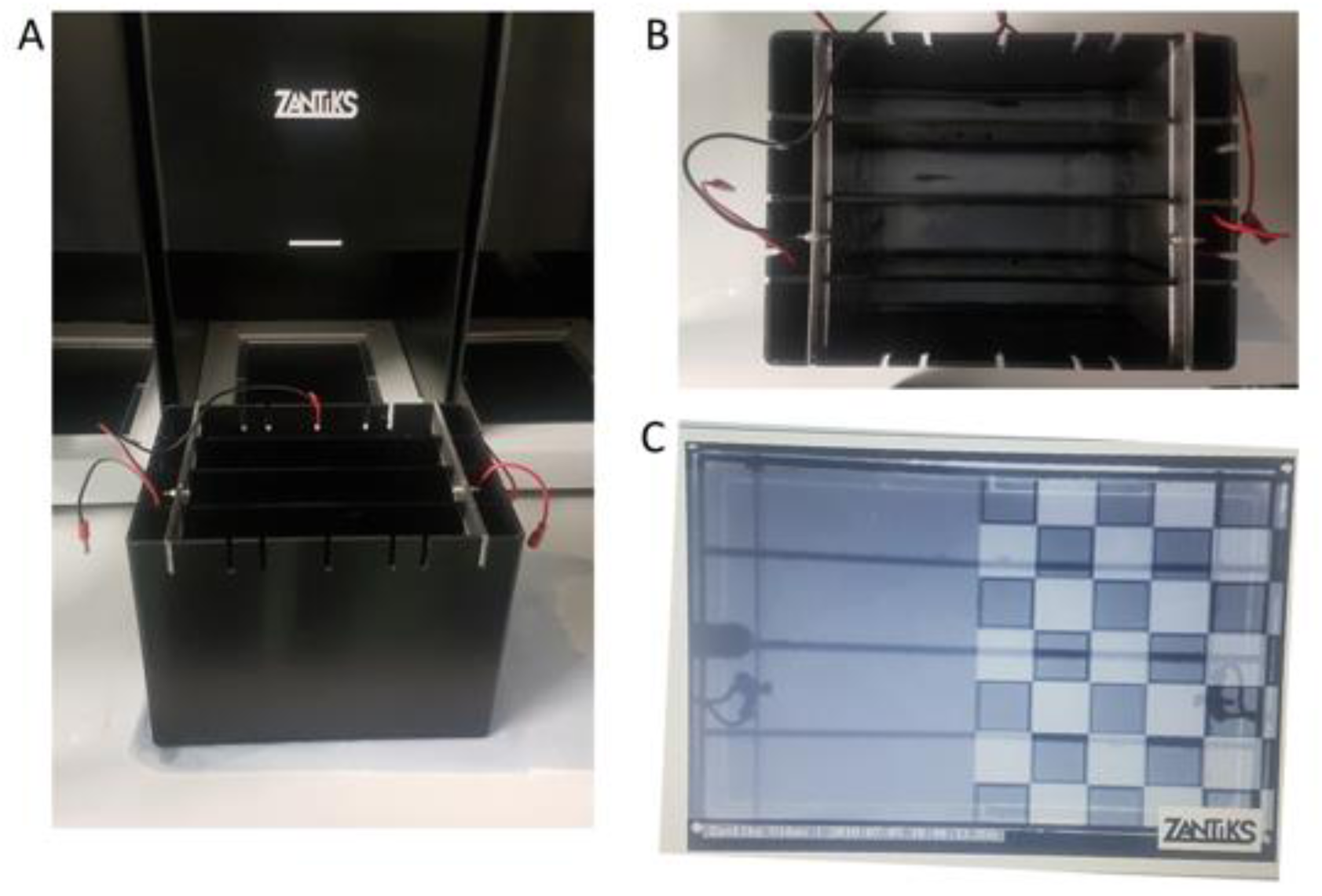
*A)*&*B)* display the tank and shocking plates used for pavlovian fear conditioning(aversive learning). *2C)*The stimuli used were based on pervious work (29), and comprised either a checker board design (‘check’)(black/white alternating squares) or a dark grey (‘grey’) background. Each tank comprised four lanes, separated by opaque acrylic dividers. At each end of the tank was located a steel plate, capable of passing a mild electric current through the tank (9v). Tanks were filled with 1L of water.

Figure 3 displays the tank set up for the y-maze alternation test. Fish were placed into one of two removable acrylic y-mazes (arm diameter × length = 70mm × 15mm) in 1L aquarium water (~40mm depth of water). Fish were filmed from above for a period of 1-hour.

**Figure 3.**
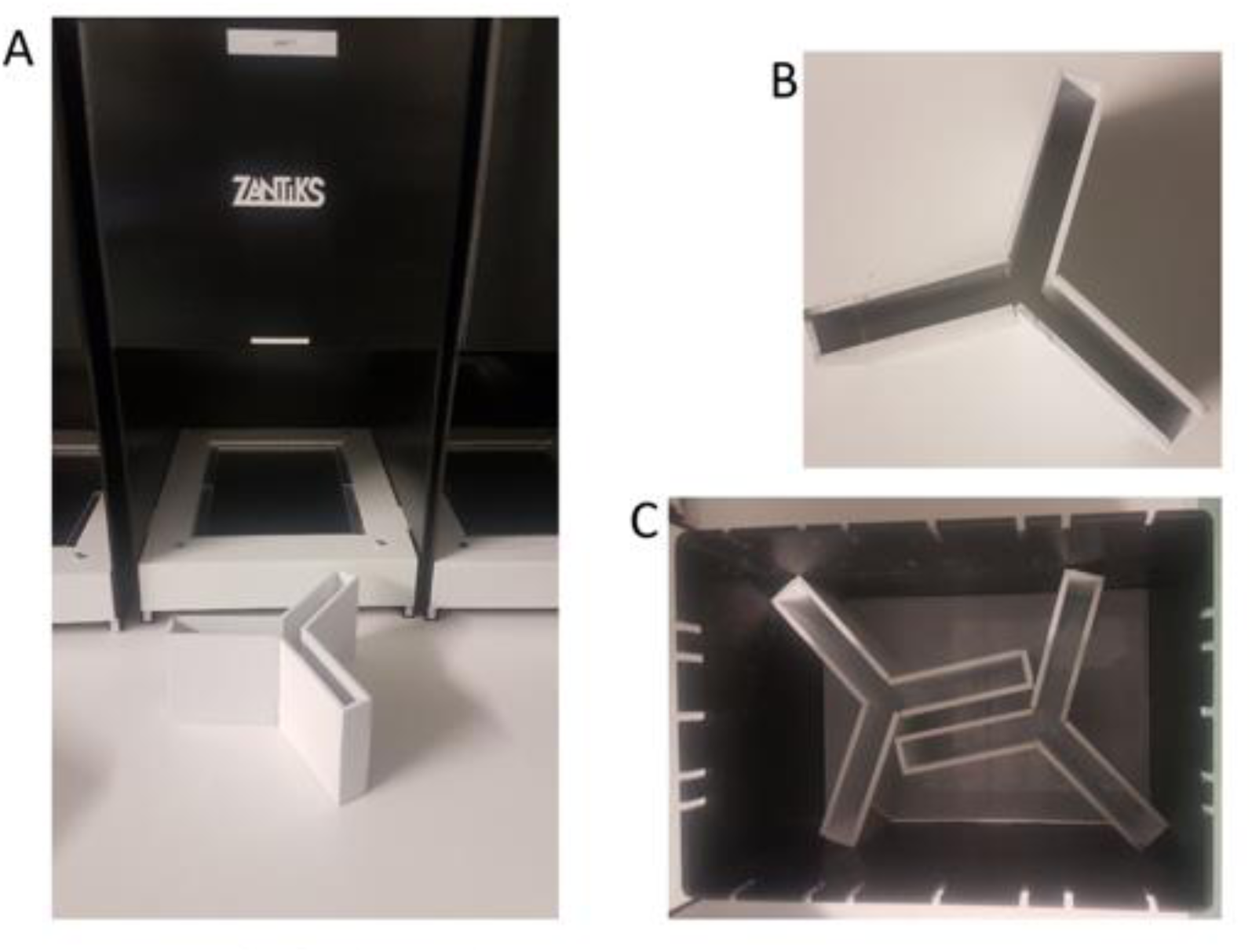
*A)*&*B) show the y-maze used to test spontaneous alternation*. *3c)* displays the tank set up for the y-maze alternation test. Fish are placed into one of two removable y-mazes in 1L aquarium water.

### Procedure

#### Appetitive conditioning

N = 16 adult (n = 8 control, n = 8 20mM ethanol) zebrafish were tested on the appetitive conditioning protocol at 3-months of age. During appetitive conditioning training, fish were housed in pairs in 2.8L tanks (divided breadth-wise with transparent mesh dividers). The training was divided into two phases, and learning was assessed following the second phase. Initially, fish were shaped for one week to associate the noise of the feed dispenser with delivery of a small amount of ZM200 (~2mg). In the second phase, fish were trained that swimming into an initiator area (Figure 1, red zone) resulted in food delivery. They were trained on this protocol for 3 days (~30-trials/day). Following completion of training, we tested learning with a series of probe trials, during which fish were exposed to 5 trials of initiator light “on” (30-sec), or “off” (30-sec), and measured total number of entries to the initiator during light on/light off.

#### Pavlovian fear conditioning

N = 44 adult (n = 22 control, n = 22 20mM ethanol) zebrafish were tested on the Pavlovian fear conditioning protocol at 3-months of age. Fear conditioning was based on a protocol developed based on published data (30). First, fish were placed, individually, into one of four lanes in each Zantiks tank. Into each tank was placed n = 2 ethanol and n = 2 control fish (position of treatment counterbalanced between trials). Initially, fish were habituated to the test environment for 30 mins, during which the base of the test tank was divided in half, lengthways, with the ‘check’ and ‘grey’ stimuli occupying half of the base each (see figure 2c), switching position every 5-min. Baseline preference was ascertained over 10-min. Following baseline preference assessment, conditioning was carried out, during which the conditioned stimulus (CS+; full screen of ‘check’ or ‘grey’, randomized between subjects) was presented for 1.5-sec, at the end of which was delivered the unconditioned stimulus (US) a brief, mild shock (9v DC, 80ms), followed by an 8.5-sec inter-trial interval (ITI), during which the non-CS (CS-) exemplar was presented at the bottom of the tank. The CS+/US was presented nine times. Following conditioning, avoidance of CS+ was ascertained by repeating the baseline, presenting both CS+ and CS-simultaneously for 1-min, and switching positions after 30-sec.

#### Y-maze test of perseveration and repetitive alternation

N = 28 adult (n = 14 control, n = 14 20mM ethanol) zebrafish were used for the y-maze test, at 3 months of age. In order to characterize perseveration and alternation in zebrafish, we designed a simple y-maze, with three identical arms. Altough memory in zebrafish using a y-maze has been previously evaluated (i.e, by blocking one arm, and exploring use of the novel arm once it is opened during training), simple unconditioned y-maze search patterns and performance has yet to be evaluated in zebrafish, and this study represents the first reports of this protocol. There were no intra-maze cues, but extra-maze (distal) cues were visible from each maze (e.g., the walls and open side of the Zantiks equipment), providing egocentric cues and allowing the fish to orient within the apparatus. The experimenter was not visible to the fish at any point during the protocol. Fish were recorded in the y-maze apparatus for up to one hour, or until they performed 100 arm-entries, whichever the sooner (here, all fish performed 100 entries in the allowed hour). This allowed for 97 overlapping series of four choices (tetragrams), of which there were a total of 16 manifestations possible. Two tetragrams (RRRR and LLLL) represented pure repetitions, and two (RLRL, LRLR) pure alternations. A completely random search strategy would be to choose every potential tetragram equally (97/16 = 6-times). However, perseverant responses sequences may encompass above-average repetitions of alternations or of repetitions. Previous research using a T-maze in which each arm was baited with an equal probability reinforcement, has demonstrated that mice tend to show generally higher levels of alternations between arms (LRLR, RLRL) than other alternatives (31).

### Data analysis

Data were analyzed in IBM SPSS^®^ Statistics for Macintosh (Version 24). Appetitive conditioning was measured by examining both acquisition data, and a series of probe trials to test learning. Acquisition data was compared between control and 20mM ethanol-treated fish using a general linear mixed model, with fixed factors as treatment (2-levels: control, 20mM ethanol) and day (3-levels), and their interaction, and ID as the random effect (to account for non—independence of repeated observations). Denominator degrees of freedom were estimated using the Satterthwaite approximation. Probe trials comprised count data, and were fitted to a generalized linear mixed effects model (Poisson distribution, log link function), with fixed effects as treatment (control, 20mM ethanol) and light status (light on, light off) and the random effect as fish ID (to account for non-independence of replicates). Denominator degrees of freedom were estimated using the Satterthwaite approximation. Fear conditioning was assessed by comparing change in preference for a stimulus following conditioning with 9 × 9v shocks. A two-way, mixed design analysis of variance (ANOVA) was applied, with proportion of CS+/CS-preference as the dependent variable, ethanol treatment as the between-subjects factor (2-levels; control, 20mM ethanol) and conditioning stage as the within-subjects factor (2-levels; pre- and post-conditioning). Finally, in order to examine perseveration in the y-maze, we carried out two analyses. In the first, we considered whether there were differences in the frequency of each of the 16 tetragrams as a function of treatment. We fitted frequency data to a generalized linear mixed effects model (Poisson distribution, log link), with treatment (control, 20mM ethanol) and tetragram (16-levels) as fixed factors, and ID as the random effect (to account for non-independence of replicates). Denominator degrees of freedom were estimated using the Satterthwaite approximation. Next, to assess whether there were any effects of developmental ethanol on frequency of pure alternations (LRLR + RLRL) or pure repetitions (RRRR + LLLL). This was assessed by fitting generalized linear models (Poisson distribution, log link function) to alternation and repetition frequency data. In both models, fixed factor for each was ethanol treatment (2-levels: control, 20mM ethanol).

## Results

### Moderate developmental ethanol exposure does not affect appetitive learning in zebrafish

Figure 4a displays the acquisition of learning data for the 20uM ethanol-treated and control fish. A linear mixed effects model revealed a significant main effect of day, *F* (2,28) = 9.15, *P* < .01 (Day 1 vs Day 2, *P* = .9; Day 1 < Day 3, *P* =.001; Day 2 < Day 3, *P* =.001), but no significant effect of treatment, *F* (1,14) = 1.17, *P* = .3, or day × treatment interaction, *F* < 1. Figure 4b displays the probe trial following appetitive conditioning, in which fish were presented with the stimulus light 5-times (10-sec), interspersed with non-light presentations (10-sec). A generalized linear mixed effects model (Poisson distribution, log link function) was fitted to the data, with number of entries to the stimulus zone as the response variable, treatment and lights-on/off as the fixed factors, and ID as the random effect. There was a main effect of lights on/off, *F* (1,18) = 9.41, *P* < 0.01 (Lights ON > Lights OFF), but not effect of ethanol treatment or lights on/off × treatment interaction, *F*s < 1.

**Figure 4.**
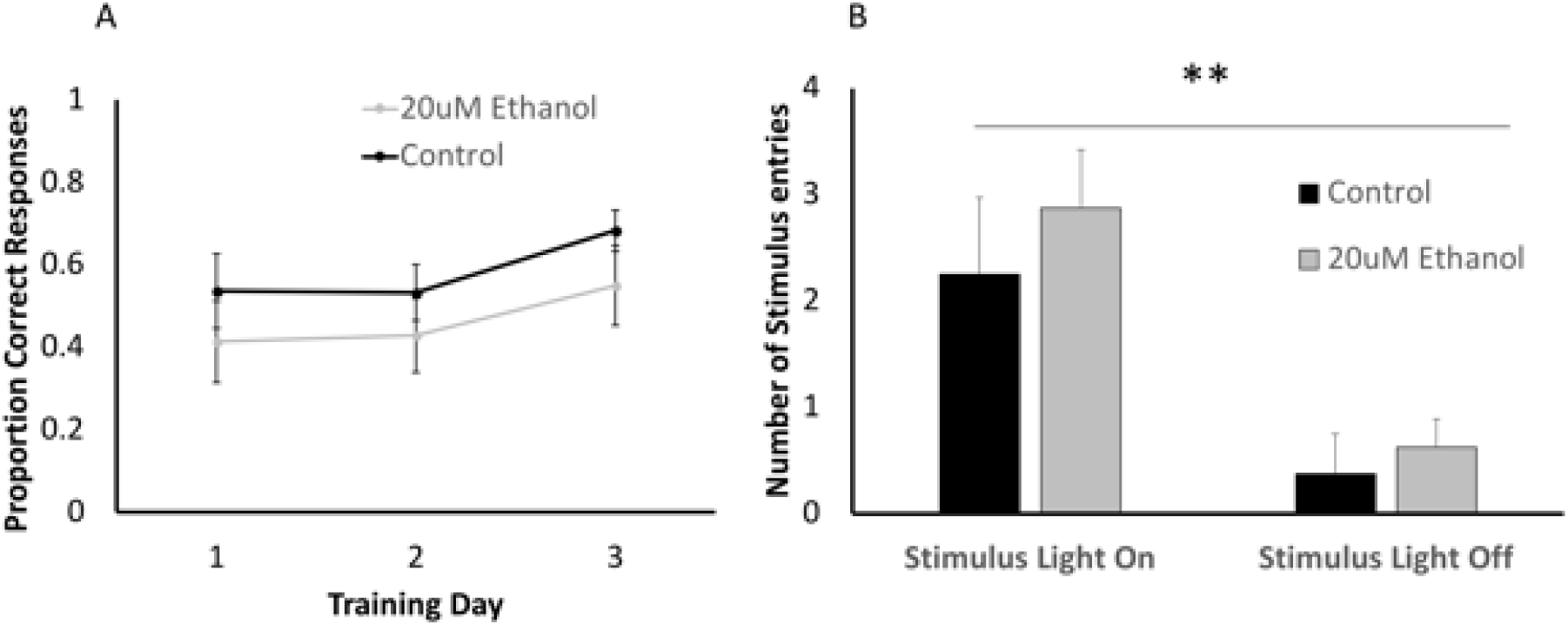
A) Training data (appetitive conditioning) for 20uM-treated and control fish. B) probe trial data (appetitive conditioning) for 20uM-treated and control fish.

### Moderate developmental ethanol exposure does not affect fear conditioning in zebrafish

Figure 5 displays the mean preference for the conditioned stimulus following Pavlovian fear conditioning, during which fish were given 9 CS + US (shock) pairings with either a checker-board, or grey image. A 2-way mixed design ANOVA revealed a significant main effect of conditioning, *F* (1,42) = 62.79, *P* < .001, with fish showing a robust reduction in preference for the conditioned stimulus. There was no main effect of ethanol treatment (*F* < 1) nor conditioning*ethanol treatment interaction (*F* < 1).

**Figure 5.**
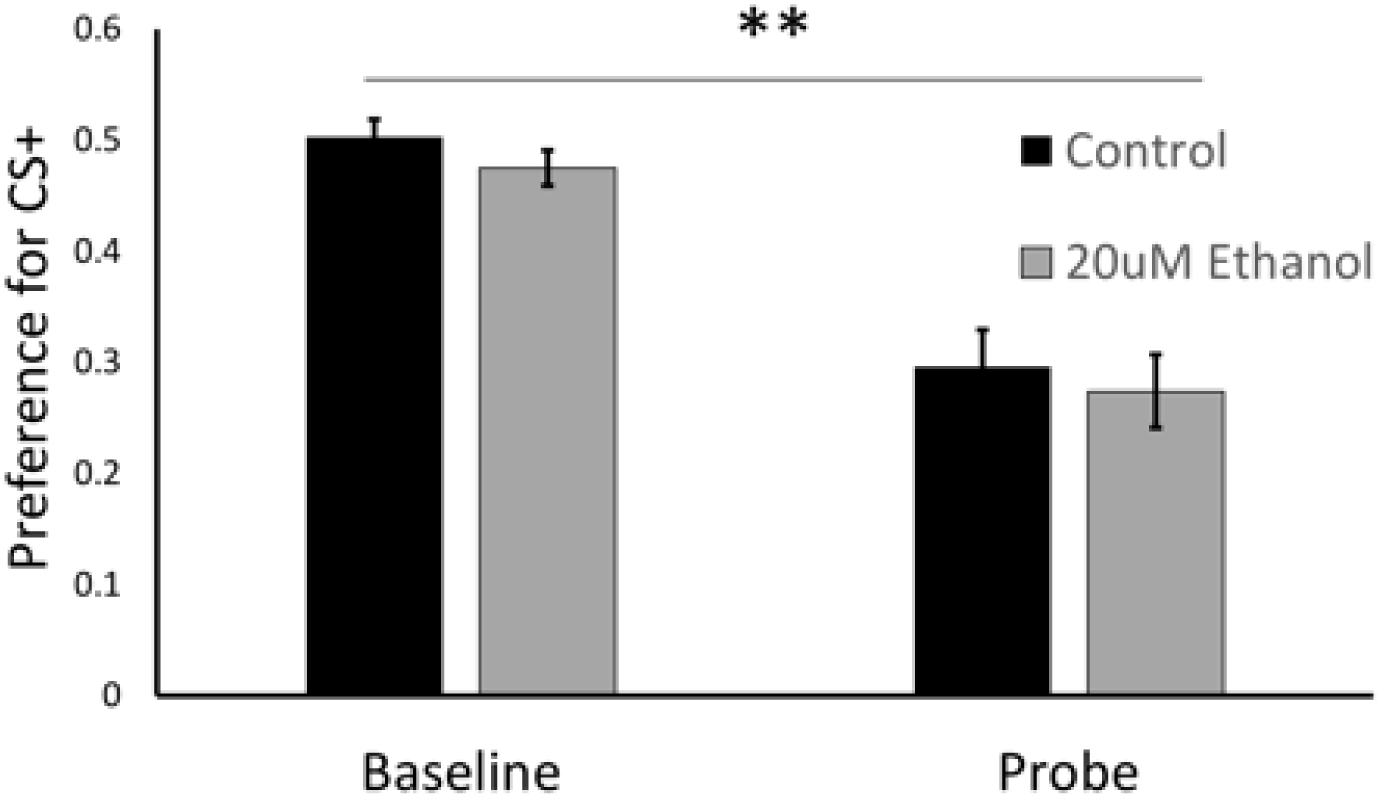
Preference for conditioned stimulus (CS+) prior to conditioning (baseline) and following pairing with 9 × 9v shocks (probe).\*\**p* < 0.0001.

### Moderate development ethanol exposure reduces alternations in a y-maze

Figure 6a displays the total choices for each tetragram during a 100-trial search period for zebrafish in a y-maze. A generalized linear mixed effects model, with Poisson distribution specified and a log link function, revealed a significant main effect for ‘tetragram’, *F* (15, 416) = 8.74, *P* < .001, characterized as significant increases in frequency of alternations (lrlr, rlrl; *P* < .001). There were no main effects of ethanol treatment (*F* < 1), nor tetragram*ethanol treatment interaction (*F* < 1). Figure 6b displays the frequency of pure alternations, and figure 6c, pure repetitions. Generalized linear models (Poisson regression) revealed a significant effect of ethanol treatment on alternations (Figure 6b; χ^2^ [df = 1] = 3.98, *P* =.046), but not on repetitions (Figure 4c; χ^2^ [df = 1] = .3, *P* = .58).

**Figure 6.**
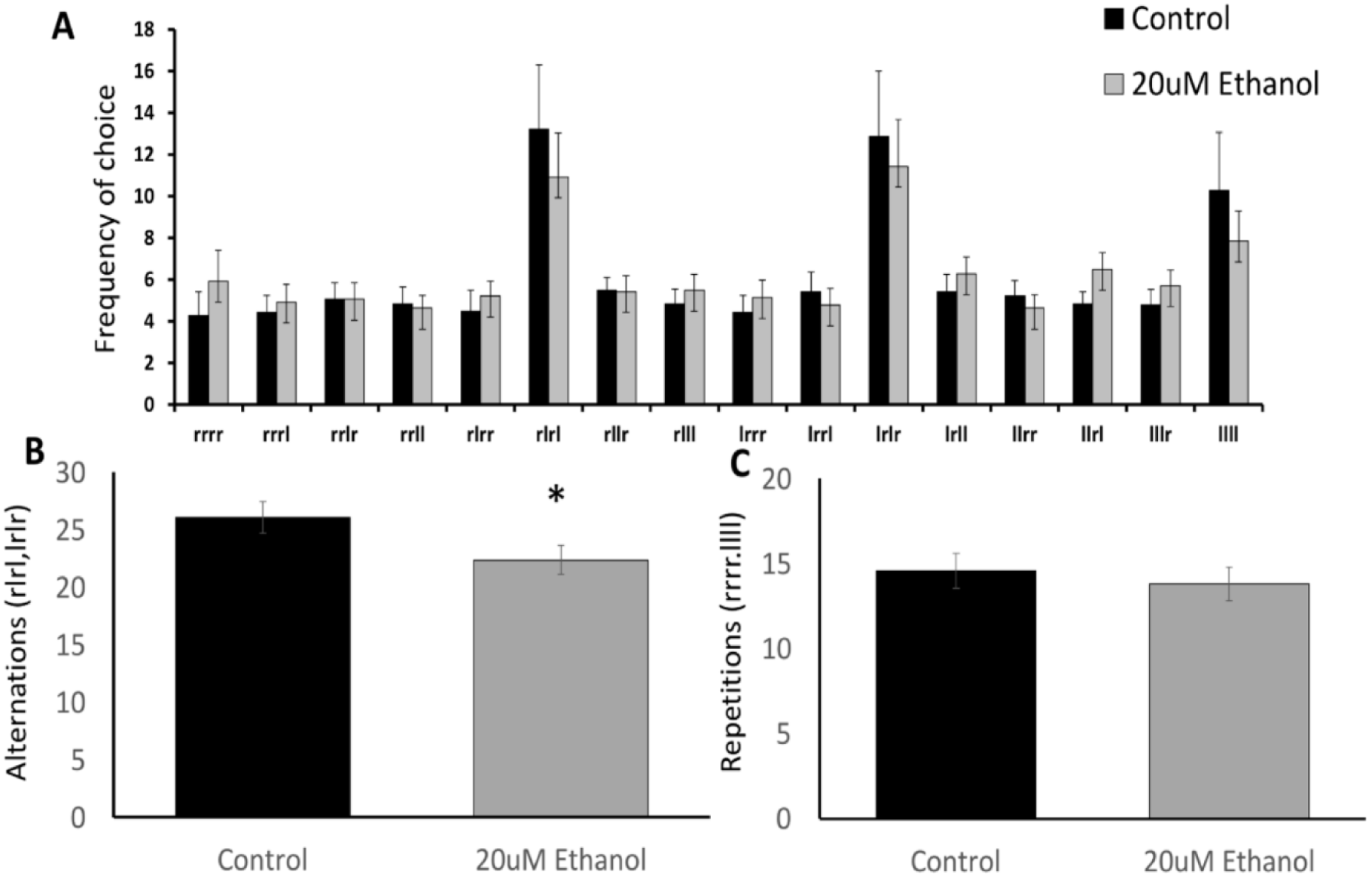
A) Frequency distribution of tetragrams during unconditional search in a y-maze. Data relate to n=14 ethanol-treated and 14 control zebrafish and comprise mean (±SEM) choices during 100 trials. B) Mean (±SEM) total pure alternations for control and 20mM ethanol-treated fish.C) Mean((±SEM) total pure repetitions for control and 20mM ethanol-treated fish.\**P* < 0.05

## Discussion

In this study we combined operant and Pavlovian learning tasks with a spatial memory task to test whether moderate levels of ethanol exposure during early brain development in zebrafish affected aspects of learning and working memory. We found that adults developmentally treated with 20mM ethanol (the equivalent of 2-3 drinks in a single sitting) performed equally well as control groups in Pavlovian and operant learning tasks, but in a spatial memory task had a decreased frequency of pure alternations (LRLR, RLRL) in a y-maze. These observations indicate that MPE can cause disruption in specific aspects of learning that persist into adulthood even when executive function is not obviously affected. This highlights the difficulty in diagnosing milder forms of FASD, but further supports the growing body of evidence implementing low to moderate levels of alcohol exposure in cognitive and behavioral abnormalities.

### Appetitive and aversive learning tasks

Operant learning tasks are used to measure executive functions involved in learning, planning, motivation and memory (30), higher functions that are often impaired in children with FASD (32,33). Here we found that exposure of larvae to 20mM ethanol [0.12% v/v] from 2-9 dpf had no detectable differences in executive function in adulthood when tested in appetitive or aversive learning tasks. This was irrespective of whether the task was under a positive (food reward) or negative (shock) stimulus control. These findings are in contrast to models of high ethanol exposure which have reported marked deficits in behavioral tasks of exposed groups (17,34,35). Although these findings may seem at odds with our own results, this disparity was not unexpected. Obvious deficits in executive function are found in humans with severe forms of FASD which are associated with periods of binge drinking during pregnancy (36). The effects of high doses of ethanol exposure have been replicated in a number of primate (37), rodent (38,39) and zebrafish models (5,18), many showing a dose-dependent effect on learning (18). Children also show a dose-dependent effect of alcohol exposure resulting in extreme variability between mild and severe cases (6). Those exposed to moderate intrauterine alcohol levels may not display obvious deficits in executive function, but have subtler effects that may not be evident until later in life. A study by Carvan III, et al (2004) looked at learning and memory in low to high doses of ethanol exposure in zebrafish and found a dose-dependent effect on learning and memory when embryos were exposed to low doses ranging from 10mM-30mM ethanol (18). The disparity between our study showing no learning deficits at 20mM could be explained by differences in the timing of doses. In their study dosing began at 4 hpf compared to 48 hpf in this study. Other studies have also found discrepancies in dose amount and effects on learning and memory tasks, a key difference between them being the timing and duration of dose (40,41). At 48 hpf zebrafish larvae are almost completely developed, the brain has developed into 5 distinct lobes, the circulatory system has developed and the heart is beating, fins develop and there is sensitivity to the environment and uncoordinated movements (42). This is the equivalent to late stages of development in the human fetus. We suspect that full brain development at the time of dosing may be a reason for executive functions still being intact, and showing no obvious signs of dysfunction.

Our findings highlight two major influences on severity and variability in FASD. Firstly, from findings of higher-dose animal models (17,18,35), we can draw the conclusion that doses higher than 0.12% v/v of ethanol exposure [up to 3% v/v] are required to cause functional deficits in executive function, specifically in goal-orientated reward tasks. Secondly, dosage is not the only key factor causing developmental defects, timing and duration of exposure are also crucial and can cause marked differences in cognitive and behavioral abilities even at lower doses of ethanol exposure (18,40). This ability to mirror the variability seen in human forms of FASD further strengthens the use of zebrafish as a model of complex neurological disorders.

### Y-maze Spontaneous Alternation Task

The y-maze has been established as a reliable protocol for testing spatial working memory in zebrafish (26). For the first time, here we use unconditioned free-search of the y-maze as a behavioral test for fish prenatally exposed to moderate levels of ethanol. We observed that all fish had a tendency for highly alternating sequences (pure alternations, LRLR or RLRL). However, 20mM ethanol-treated fish used pure alternation as a search strategy significantly less than their untreated counterparts. Using tetragram configurations, similar to those used by Gross, et al (2011) (43), allowed us to compare random or specific search strategies employed by fish when swimming freely in a y-maze. We would expect that in a reward-absent maze, with no highly salient intra or extra-maze cues, that fish would choose each search configuration equally (i.e. in 100 trials each tetragram should be chosen n=6 times). However, for both treated and nontreated groups this was not the case. Our findings that all subjects have an increased preference for pure alternations was previously seen by Gross, et al (2011) (43) with rats and Neuringer (1992) (44) with pigeons. Both studies similarly reinforced each option equally (here this was done by an absence of reward, opposed to associating each arm with a reward an equal number of times). This observation could be explained by the ‘*law of least mental work*’, an adaptation from Hull’s ‘*law of less work*’(45), which suggests that in a non-reinforced task subjects will opt for the behavior that is least cognitively demanding. Simplified strategies, like repeating the last action, may be favored in non-discrete behavioral tasks, possibly identifying repetition of pure alternations as the least cognitively demanding strategy. The random incorporation of the other 14 possible search strategies could be influenced by information seeking or a desire for change or novelty (46), but in the absence of any new information being presented the organism goes back to the least mentally demanding search pattern.

The reduction of pure alternations in ethanol treated fish is contradictory to what we would expect. Several animal models have used a form of spontaneous alternation to measure working memory, whether it is in a t-maze (43,47), y-maze (48) or plus maze (49), or the use of two choice guessing tasks in humans (50). Over a range of conditions frequency of pure alternations has normally been reported as higher in the treated group than in the control groups (43,50). However, the mechanisms affected by moderate ethanol exposure that result in the deficit that we see here, are not completely clear. It is possible that these differences are due to damage to the hippocampus, effecting working memory. Rats exposed to ethanol within the first 2 weeks of neonatal life, the rodent equivalent of the third trimester in humans, have been reported to have functional impairments of hippocampal neurons, specifically in the CA3 region (51–53). This could potentially explain deficits seen here, as rodents with hippocampal lesions also perform poorly in spatial memory tasks, specifically with familiarization (25). However, if there were hippocampal impairments, we would expect to see some evidence of this during either of the other two operant learning tasks used. Even though these trials require extensive training and rely on associative memory to be successfully performed, they also require the hippocampus, and during the early process of learning, spatial working memory, short-term memory and the ability to convert experiences into long-term memories. The combination of learning and memory tasks used and the paralleled ability of treated and control groups in the operant learning tasks thus suggest that memory may not be a fault for the changed behavioral pattern seen in the y-maze by treated fish.

An alternative explanation is the theory of “choice hysteresis”. Choice hysteresis is the tendency of animals and humans to repeat recent choices. Bonaiuto, de Berker and Bestmann (2016) (54) describe a virtual model in which activity decay leftover from recently activated neural circuits increased repetition biases. They also describe how depolarizing or hyperpolarizing the dorsolateral prefrontal cortex (dlPFC) could increase or suppress choice hysteresis, respectively. Children born with FASD are often reported to have deficits in the dlPFC as is established by the Wisconsin Card Sorting task (neuropsychological test for executive function which is used to determine damage to the dlPFC) (55). Again, this is another area in which severity and variability are in strong correlation with dose of alcohol exposure, with high, chronic doses of alcohol exposure, (i.e. those born to alcoholic mothers) performed the poorest on this task compared to those exposed to lower doses of alcohol and control groups.

This is the first time that the y-maze has been used in the context of FASD and there are evident differences in the search strategies employed by treated and nontreated groups. However, the lack of comparative studies carried out at other doses limits what can be inferred from this data and therefore mechanisms and meanings are limited to theoretical ideas. Although many other studies have investigated the effects of PNEE on spatial working memory (7,56–58), the lack of standardized measures of spontaneous alternation makes it difficult to compare findings from one study to another unless they have used similar data analysis. Also, significant differences can be seen between rewarded and reward-absent tasks. From this study we can concluded that moderate PNEE does alter behavior in a spontaneous alternation task. However, at this stage we can only hypothesize about possible mechanisms responsible for this change. It is also difficult to judge how this may relate to human behavior, thus, further work would be required to draw out anything conclusive.

### Limitations and future directions

There are limitations to this study. First, only one dose and one exposure time were used. In light of differences on learning and memory seen in other behavioral studies (see review: (13)) future studies could use variations of these two factors to test if there are any behavioral changes in the tasks performed here. Most interestingly would be the effect on spontaneous alternations in the y-maze. We predict that exposure to high doses of ethanol, to the extent that it causes cognitive impairment, would cause an increase in pure repetitions (LLLL,RRRR) and pure alternations would occur at a rate equal to that of controls. Secondly, we used a mixture of male and female fish for our experiments. Many studies have found effects of sex in some species following developmental ethanol exposure (57,59). This may represent a limitation of the fish model in terms of translational relevance, as it appears at odds with other vertebrates in this regard. Finally, our limited repertoire of behavioral tasks may result in other behavioral abnormalities being missed. Future studies should incorporate more tasks involving complex cognitive functions to fully elucidate the effect of moderate PNEE.

### Concluding remarks

In regard to the primary aim of this study we found that MPE caused marked behavioral changes in the search strategy employed by treated fish in a y-maze. However, the dose and timing of exposure did not impact on executive functions required for Pavlovian operant learning tasks. Thus, we can conclude that even in the absence of physical malformations, developmental exposure to moderate levels of ethanol can cause subtle behavioral and cognitive changes that persist into adulthood.

Despite decades of investigation and a mountain of data examining the toxic and deleterious effects of alcohol on the developing fetus, the number of children born with FASD is on the rise (60). Being the leading form of preventable mental retardation and having lifelong effects that can severely reduce the quality of life of those affected (61), it is critical that we fully understand the impact that all levels of ethanol exposure can have. The effect of moderate levels of alcohol exposure are becoming even more crucial with the number of women having unplanned births and pregnancies being as high as 23% and 40% respectively (62,63), worldwide. Therefore, the chance of drinking before becoming aware of being pregnant is a huge risk factor. Doses as low as 10mM have been reported to affect cognitive abilities, this is the equivalent of 1-2 drinks in one sitting (18). This is also the current UK recommend ‘safe level’ of consumption during pregnancy (64). With guidelines like these in place and a growing body of evidence suggesting detrimental effects of low and moderate levels of alcohol exposure, research in these areas is becoming even more important to help change health advice and the way society see maternal drinking.

## Acknowledgements

This work was funded by a University of Portsmouth Science Faculty PhD studentship (MC).

